# Canonical Wnt signaling regulates patterning, differentiation and nucleogenesis in mouse hypothalamus and prethalamus

**DOI:** 10.1101/355750

**Authors:** Elizabeth A. Newman, Dan Wu, Makoto Mark Taketo, Jiangyang Zhang, Seth Blackshaw

## Abstract

The hypothalamus is a small, but anatomically and functionally complex, region of the brain whose development is poorly understood. In this study, we have explored its development by studying the canonical Wnt signalling pathway, generating gain and loss of function mutations of betacaten in(*Ctnnb1*) in both hypothalamic and prethalamic neuroepithelium. Deletion of *Ctnnb1* resulted in an anteriorized and hypoplastic hypothalamus. Posterior structures were lost or reduced, and anterior structures were expanded. In contrast, overexpression of a constitutively active mutant form of *Ctnnb1* resulted in severe hyperplasia of prethalamus and hypothalamus, and expanded expression of a subset of posterior and premamillary hypothalamic markers. Moderate defects in differentiation of *Arx*-positive GABAergic neural precursors were observed in both prethalamus and hypothalamus of *Ctnnb1* loss of function mutants, while in gain of function mutants, their differentiation was completely suppressed, although markers of prethalamic progenitors were preserved. Multiple other region-specific markers, including several specific posterior hypothalamic structures, were also suppressed in *Ctnnb1* gain of function mutations. Severe, region-specific defects in hypothalamic nucleogenesis were also observed in both gain and loss of function mutations of *Ctnnb1*. Finally, both gain and loss of function of *Ctnnb1* also produced severe, cell nonautonomous disruptions of pituitary development. These findings demonstrate acentral and multifaceted role for canonical Wnt signalling in regulating growth, patterning, differentiation and nucleogenesis in multiple diencephalic regions.

**Highlights:**

- Canonical Wnt signalling regulates anteroposterior patterning in the hypothalamus.
- Canonical Wnt signalling regulates differentiation of GABAergic neurons in both prethalamus and hypothalamus.
- Canonical Wnt signalling regulates differentiation and nucleogenesis of multiple hypothalamic neuronal subtypes.
- Canonical Wnt signalling in hypothalamic neuroepithelium regulates pituitary morphogenesis and differentiation.

## 1. Introduction

The hypothalamus regulates a broad range of homeostatic processes and innate behaviors (Bedont et al., 2014; Burbridge et al., 2016; Caron and Richard, 2017; Herrera et al., 2017; Lechan and Toni, 2000; Lee et al., 2012; Liu et al., 2017; Morrison, 2016). In spite of its great physiological and behavior importance, the development of the hypothalamus is poorly understood, in large part because of its great anatomic and cellular complexity (Braak & Braak, 1992; Flament-Durand, 1980; Lechan & Toni, 2000). Although recently some progress has been made in identifying extrinsic and intrinsic factors that control hypothalamic patterning and neurogenesis, much remains unknown (Bedont et al., 2015; Burbridge et al., 2016; Xie and Dorsky, 2017). In particular, we understand little about how classic growth and differentiation factors regulate spatial patterning of hypothalamic neuroepithelium.

In contrast to the cerebral cortex (Fukuchi-Shimogori and Grove, 2001, 2003; Toyoda et al., 2010) and thalamus (Kiecker and Lumsden, 2004; Scholpp et al., 2006; Vieira et al., 2005), where signaling centers produce spatial gradients of differentiation factors to pattern these areas, such signaling centers have yet to be discovered in the hypothalamus. While several studies have demonstrated the necessity of Shh (Shimogori et al., 2010; Szabo et al., 2009), BMP (Manning et al., 2006; Ohyama et al., 2008), and FGF signaling (Bosco et al., 2013; Pearson et al., 2011) in specification and/or survival of specific hypothalamic cell types, a defined role for these factors in hypothalamic patterning has not been identified.

An additional factor complicating studies of hypothalamic patterning is the lack of consensus about the orientation of the hypothalamus relative to other forebrain structures such as the prethalamus and telencephalon. The prosomere model of forebrain development places the hypothalamus anterior to the prethalamus and ventral to the telencephalon (Ferran et al., 2015; Puelles and Rubenstein, 2015), while columnar models place it ventral to the prethalamus and posterior to the telencephalon (Swanson, 1992, 2012). These models are generated by differing interpretations of developmental gene expression patterns, and disproving either one requires a better understanding of the genes that pattern the hypothalamus and associated structures.

Canonical Wnt signaling is essential for anteroposterior patterning of the vertebrate forebrain (Kiecker and Niehrs, 2001) and is an excellent candidate mechanism for regulating patterning of both the hypothalamus and prethalamus. Multiple sites of Wnt expression are found along the margins of both structures. Prior to the onset of neurogenesis in the mouse, both *Wnt3/3a* and *Wnt8a/b* are produced by cells in the thalamus adjacent to the zona limitans intrathalamica (ZLI) and hypothalamus, respectively (Fig. 1a) (Braun et al., 2003; Martinez-Ferre et al., 2013). This expression pattern is maintained following the onset of neurogenesis (Fig. 1b), with the addition of a minor anterior zone of Wnt8b expression at the hypothalamic/telencephalic border (Fig. 1b), and a broad zone of Wnt7a/b expression throughout the prethalamus and in progenitors of GABAergic neurons in the hypothalamus (Shimogori et al., 2010).

**Figure 1:**
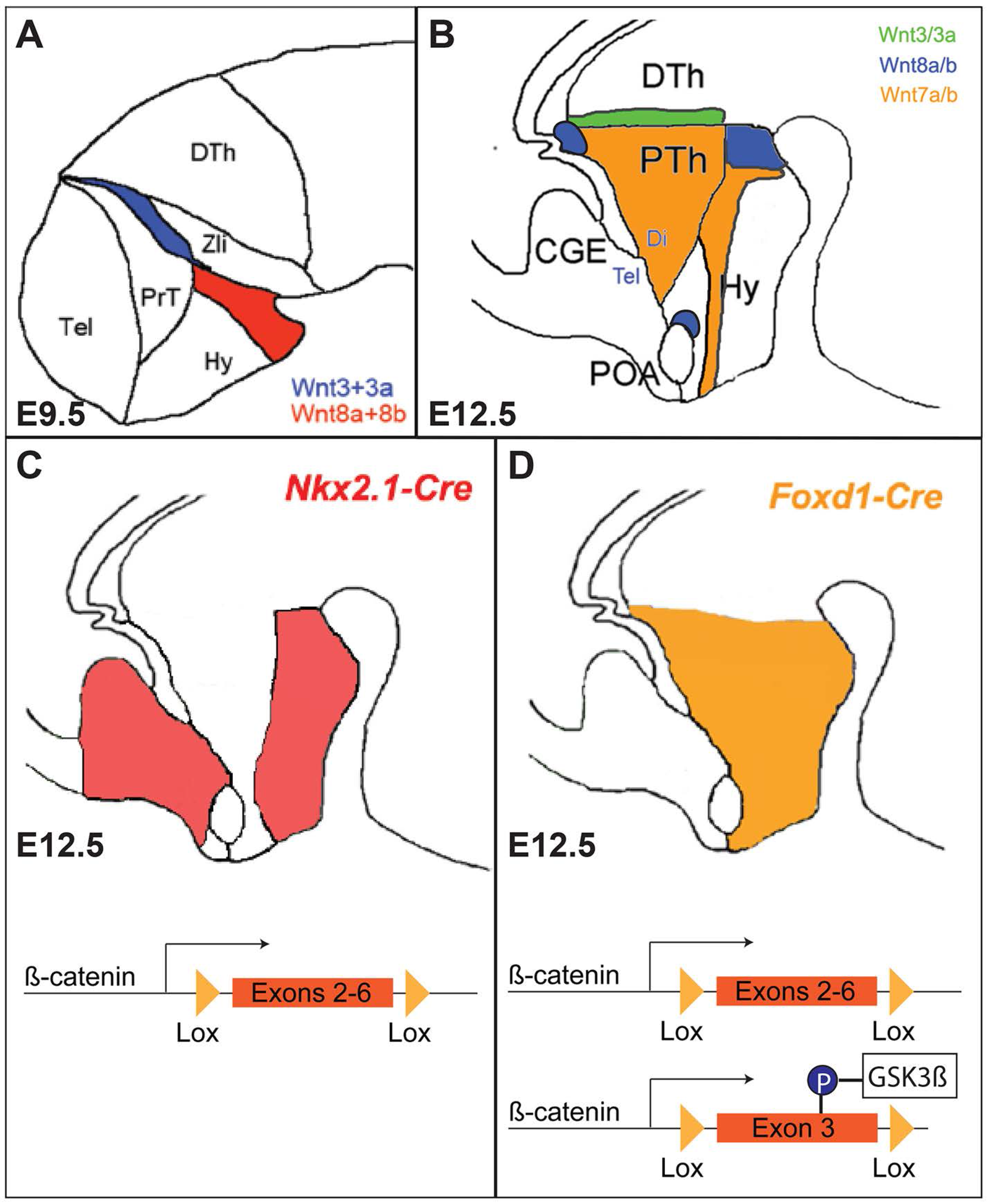
Wnt expression and Cre activity in the developing hypothalamus. A. Wnt3/3a and Wnt8a/8b are expressed in a contiguous stripe across the developing prethalamus and hypothalamus at E9.5. B. At E12.5, Wnt3/3a are expressed in a stripe across the posterior prethalamus and Wnt8a/8b is restricted to the mamillary of the hypothalamus, as well as small domains in the dorsal anterior hypothalamus and thalamic eminence. At this time, Wnt7a/7b is expressed in interneuron progenitors across the hypothalamus. C-D. Schematics showing activity patterns of *Nkx2.1-Cre* (C) and *Foxd1-Cre* (D) at E12.5. *Nkx2.1-Cre* is restricted to a ventral stripe across the anterior-posterior axis and is excluded from the prethalamus. *Foxd1-Cre* is expressed throughout the whole of the hypothalamus and prethalamus. We used a conditional deletion strategy to either excise a functionally required portion of the *Ctnnb1* gene to generate a *Ctnnb1*-null mouse, or excise an autoinhibitory domain within the third exon to generate a constitutively active form of Ctnnb1.

Previous work has shown that canonical Wnt signaling induces expression of markers of sensory thalamus while suppressing prethalamic markers (Bluske et al., 2012; Braun et al., 2003), but has not examined its role in later stages of prethalamic development. Studies of the role of the canonical Wnt pathway factors in hypothalamic patterning have primarily been conducted in zebrafish (Lee et al., 2006; McPherson et al., 2016; Russek-Blum et al., 2008; Xie et al., 2017). These studies demonstrated an essential role for *Wnt8b* and its effector gene *Lef1* in regulation of both neurogenesis and differentiation of selected hypothalamic neuronal subtypes, but did not observe an effect on hypothalamic patterning per se. None of these studies, however, studied the global effects of manipulating canonical Wnt signaling by targeting the major signaling component *Ctnnb1*. In this study, we analyze mice in which we have generated selective gain and loss of function mutations of *Ctnnb1* in hypothalamic and prethalamic neuroepithelium.

## 2 Materials and methods

### 2.1 Mouse breeding and embryo collection

Heterozygous *Foxd1-Cre: GFP* knock-in mice (*Foxd1-Cre*/+)(Humphreys et al., 2008) were mated to *Ctnnb1*^*lox/lox*^ (Brault et al., 2001) to generate *Foxd1-Cre*/+; *Ctnnb1*^*lox*/+^ mice which were then crossed back to *Ctnnb1*^*lox/lox*^ mice to generate *Ctnnb1*-null mice. The *Foxd1-Cre*/+ mice were also crossed to *Ctnnb1*^*ex3/ex3*^(Harada et al., 1999) mice to generate *Ctnnb1* gain of function mutants, by excising an autoinhibitory domain within the *Ctnnb1* gene. Mice were maintained and euthanized according to protocols approved by Johns Hopkins Institutional Animal Care and Use Committee.

### 2.2 *In situ* hybridization

Embryos were either embedded fresh frozen in OCT, or post-fixed in 4% paraformaldehyde in 1×PBS then embedded in gelatin, before sectioning. *In situ* hybridization on fresh frozen tissue was performed as previously described (Blackshaw et al., 2004). *In situ* hybridization on post-fixed sections was performed as previously described (Shimogori et al., 2010).

### 2.3 EdU staining and cell counting

Pregnant dams were given a 2 hr pulse of 50 mg/kg EdU (20 mg/mL in saline) on embryonic day E12.5 and then sacrificed via cervical dislocation. Embryos were post-fixed in 4% PFA, then cryopreserved in 30% sucrose, before being embedded in OCT and cryostat sectioned. The Click-IT EdU imaging kit from Invitrogen was used to perform the EdU staining. Slides were coverslipped using Vectashield before being imaged at 40x on a Keyence microscope. Cell counting was conducted using ImageJ on five to six sections per animal. All counting was done blinded and manually.

### 2.4 TUNEL staining

Post-fixed and cryopreserved E12.5 embryos were sectioned and dry mounted. TUNEL staining was performed according to the protocol for cryopreserved tissue in the *In Situ* Cell Death Detection Kit-TMR red (Roche). A positive control was used (3000 U/mL DNAse I). Slides were DAPI stained then coverslipped using Vectashield.

### 2.5 Statistical methods used

For the EdU cell counting, the number of EdU-positive cells was taken as a fraction of the total area of the hypothalamus measured in µm^2^. The values of the sections were averaged for each individual, and then compared using a two-tailed t-test. Three animals were used for each genotype.

### 2.6 MRI

MRI was performed on E12.5 control and beta-catenin gain of function mutant embryos. MRI data were acquired on a Bruker 11.7T spectrometer, using a 10 mm volume transceiver coil. High angular resolution diffusion MRI data was acquired using a diffusion-weighted gradient and spin echo sequence (Aggarwal et al., 2010) with the following parameters: field of view = 6×7×7.5mm, 35um isotropic resolution, 4 non-diffusion weighted images and diffusion weighted images acquired along 30 diffusion directions, b-value = 1300 s/mm^2^, 2 signal averages, echo time / repetition time = 35/600 ms, and scan time of 34 hrs. Tract density images (Farquharson et al., 2016) were generated using the MRtrix tool (www.mrtrix.org) from the diffusion MRI data.

## 3. Results

### 3.1 Loss of function of*Ctnnb1* in Nkx2.1-positive posteroventral hypothalamus disrupts Lef1 expression and leads to a reduction in posterior hypothalamic markers

To investigate the role of canonical Wnt signaling in hypothalamic and prethalamic development in mouse, we used two different genetic strategies to selectively delete *Cttnb1*. To selectively target posteroventral hypothalamus, we used the *Nkx2.1-Cre* transgene, which is active in this region from E9.5 onward (Shimogori et al., 2010; Xu et al., 2008), but is not active in prethalamus (Fig. 1c). To target both hypothalamic and prethalamic neuroepithelium, we used *Foxd1-Cre* knock-in mice, in which Cre is active in these areas beginning at E8.0 (Fig. 1d) (Newman et al., 2018). These were then crossed to floxed *Ctnnb1* mice (Brault et al., 2001) to generate *Nkx2.1-Cre;Ctnnb1*^*lox/lox*^ and *Foxd1-Cre;Ctnnb1*^*lox/lox*^ animals.

Examining *Nkx2.1-Cre;Ctnnb1*^*lox/lox*^ mice at E12.5, we observed a substantial reduction in the size of the domains expressing markers of posterior hypothalamus. These include *Foxa1* and *Irx5*, which mark supramammilary (SMM) neuroepithelium (Fig. 2a-d (black arrowhead), Fig. 2u), as well as *Foxb1* and *Sim1*, which mark the mamillary (MM) region (Fig. 2e-h black arrowhead, Fig. 2u, Fig. S1A-J) (Shimogori et al., 2010). *Sim1* is also expressed in the hypothalamic paraventricular nucleus (PvN) and the zona limitans intrathalamica (ZLI), both of which are *Nkx2.1*-negative. However, we did not observe altered expression of *Sim1* in either the PvN (Fig. 2h open arrowhead, Fig. S1A-J) or the ZLI (Fig. 2h, red arrowhead; Fig. S1A-J). The expression of *Nkx2.1* itself was reduced (Fig. S1K-U).

**Figure 2:**
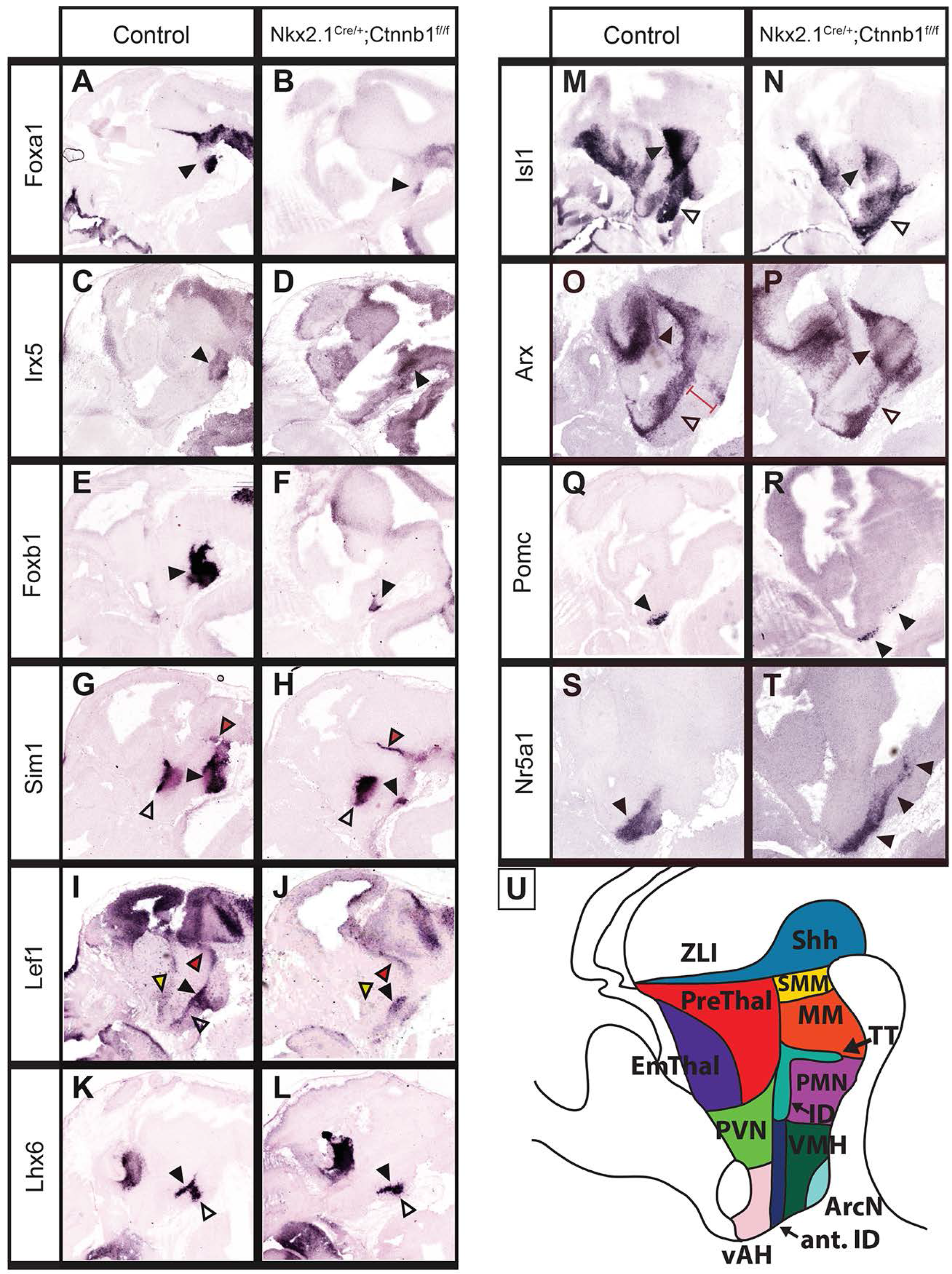
Loss of function of beta-catenin results in an anteriorization of the hypothalamus. A-T: *In situ* hybridization of hypothalamic markers on E12.5 control (*Cre*-negative, either *Ctnnb1*^*lox/lox*^ or *Ctnnb1*^*lox*/+^) and *Nkx2.1-Cre;Ctnnb1*^*lox/lox*^ sagittal sections. A-J: Black arrowheads indicate the supramamillary (A-D), mamillary (E-H) and premamillary (I-J), all of which were reduced in the mutant. The PVN was unaffected (open arrowheads, G and H). The ID lost *Lef1* expression (open arrowheads, I and J) but not *Lhx6* expression (black arrowheads, K and L). The TT was reduced (open arrowheads, K and L). M-P: Black arrowheads indicate the prethalamus, which was intact in mutants. Open arrowheads indicate the hypothalamic stripe domain, which was ventralized in the mutants. Red bar in O represents the *Arx*-negative hypothalamic domain that was missing in the mutant. Q-T: Anterior structures were expanded into the posterior domain. Q-R: black arrowheads indicate the arcuate. S-T: black arrowheads indicate the VMH. U: Schematic showing wild-type anatomy of the hypothalamus and prethalamus.

The Wnt effector gene *Lef1* is selectively expressed in premamillary neuroepithelium (PMN), in the posterior prethalamus adjacent to the ZLI (Fig. 2i-j, yellow arrowhead) (Shimogori et al., 2010), the tuberomamillary terminal (TT), (Fig. 2i-j, black arrowhead, Fig. 2u) and in the intrahypothalamic diagonal (ID) (Fig. 2i-j, open arrowhead), a small domain along the hypothalamic-telencephalic boundary the may correspond to the domain of *Wnt8b* expression (Fig.1B, Fig. 2i-j, red arrowhead, Fig. 2u). We observed that the majority of *Lef1* expression was lost, including the ID, most of the PMN and TT, and the anterior hypothalamic expression domain, although the *Nkx2.1*-negative prethalamic domain is preserved. Another marker of the ID and TT, *Lhx6*, was maintained in the ID (Fig. 2k-l, S2A-H, black arrowhead) but greatly reduced in the TT (open arrowhead). *Isl1* and *Arx*, which are expressed in contiguous domains that span the prethalamus and hypothalamus, did not display any general change in the shape or size of their expression domains in either structure (Fig. 2m-o), but were both displaced ventrally. This is most likely a result of a reduction in the *Nkx2.1*-positive, *Arx* and *Isl1*-negative hypothalamic domain of the ventral hypothalamus, which is supported by the observed reduction in the size of the *Nkx2.1* expression domain here (Fig. S1K-U).

The most anterior region of the *Nkx2.1*-postive hypothalamic domain includes the arcuate nucleus (ArcN) and ventromedial hypothalamic nucleus (VMH) (Xu et al., 2008)(Shimogori et al., 2010,). We observed that expression of both the ArcN marker *Pomc* and the VMH marker *Nr5a1* was extended posteriorly in *Nkx2.1-Cre;Ctnnb1*^*lox/lox*^ mice (Fig. 2Q-U, Fig. S2I-B’). Since the overall size of the *Nkx2.1*-positive hypothalamic region was reduced in these animals, it is unclear whether this expanded expression domain represents a change in cell fate or simply a posterior migration of *Pomc* and *Nr5a1*-positive cells.

### 3.2 Loss of function of*Ctnnb1* in*Foxd1*-positive prethalamic and hypothalamic neuroepithelium leads to a reduction in posterior hypothalamic markers, and modest reduction in prethalamic markers

Using the *Nkx2.1-Cre* line enables us to investigate the role of canonical Wnt signaling in regulating the patterning and differentiation of the posteroventral hypothalamus, but does not let us study the patterning of the hypothalamus and prethalamus more broadly. In particular, it does not allow us to determine whether canonical Wnt signaling coordinately regulates expression of genes expressed in contiguous domains across both structures. To study this, we used the *Foxd1-CreGFP* knock-in line (Humphreys et al., 2010), which we have previously used to generate selective loss of function mutations in hypothalamus and prethalamus (Newman et al., 2018; Salvatierra et al., 2014).

When we compared *Foxd1-Cre;Ctnnb1*^*lox/lox*^ mice to control littermates at E12.5, we observed a reduction in the size of both the prethalamic (black arrowhead) and hypothalamic expression domains (open arrowhead) of *Arx* and *Isl1* (Fig. 3A-D, Fig. 3U, Fig. S3A-L). We observed a near total loss of prethalamic *Lef1* expression with the exception of a portion adjacent to the ZLI (black arrowhead), along with a reduction in hypothalamic *Lef1* expression (open, red and yellow arrowheads) that closely resembles that seen in *Nkx2.1-Cre;Ctnnb1*^*lox/lox*^ mice (Fig. 3E-F, U). *Lhx6* displayed a more severe reduction in expression than in *Nkx2.1-Cre;Ctnnb1*^*lox/lox*^ mice, with only low levels detected in the ID (open arrowhead) and none in the TT (black arrowhead) (Fig. 3G-H, U). *Lhx9*, which is transcribed in precursors of hypocretin neurons in the lateral hypothalamus (black arrowheads), and the thalamic eminence (EmThal) (open arrowheads) (Dalal et al., 2013; Liu et al., 2015; Shimogori et al., 2010), showed reduced and displaced hypothalamic expression but a modestly expanded expression domain in the EmThal (Fig. 3I-j, Fig. 3U, Fig. S3M-R).

**Figure 3:**
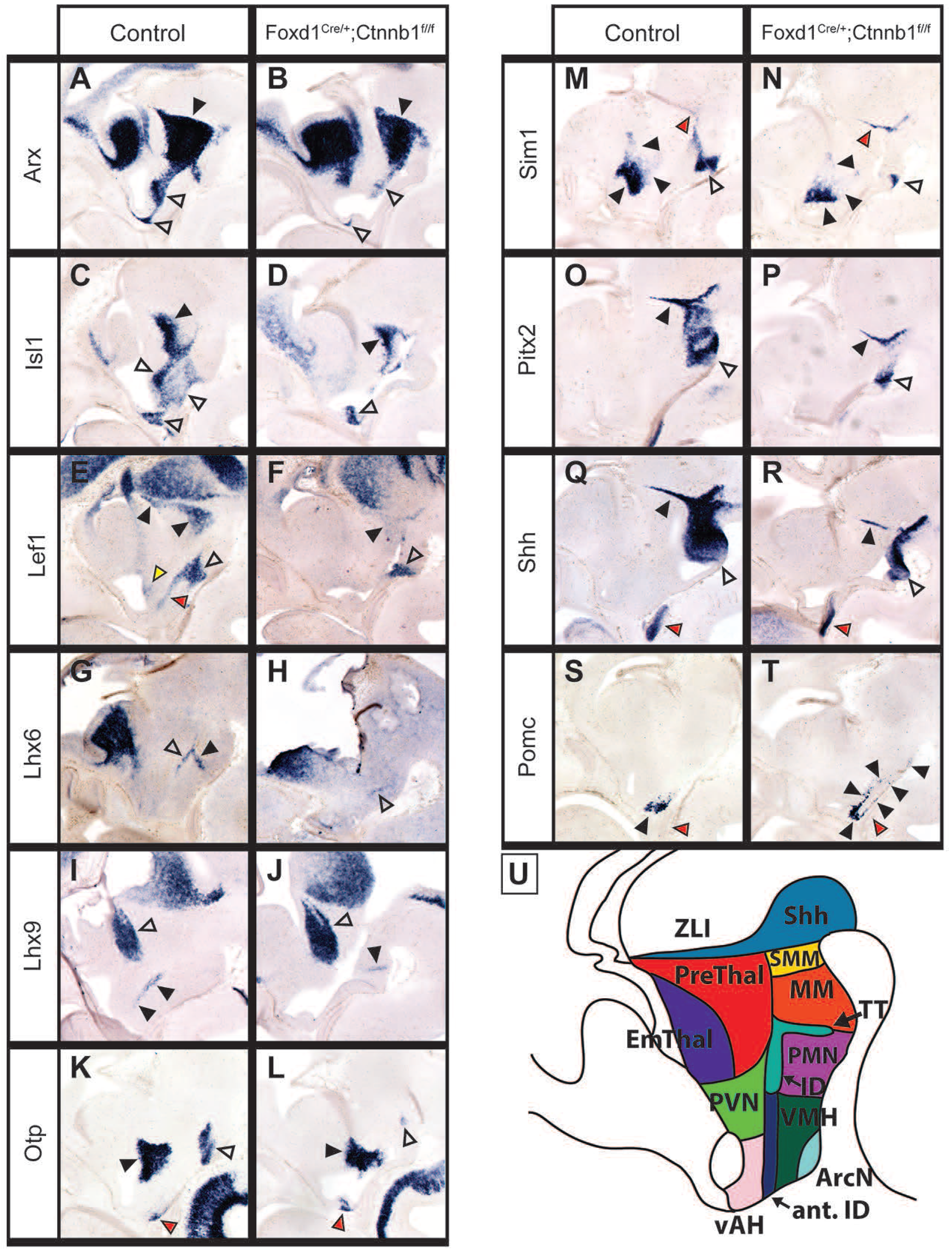
Loss of function of beta-catenin results in a thinning and anteriorization of the hypothalamus but has no effect on the prethalamus: Sagittal. A-T: *In situ* hybridization on E12.5 control (*Cre*-negative, either *Ctnnb1*^*lox/lox*^ or *Ctnnb1*^*lox*/+^) and *Foxd1-Cre;Ctnnb1*^*lox/lox*^ sagittal sections. A-D: Black arrowheads indicate the prethalamus, which was unaffected in the mutants. White arrowheads indicate the thinned hypothalamic domain. The *Lef1* domain of the ID was absent in the mutants (red arrowhead, E and F) but the *Lhx6* domain was unaffected (open arrowhead, G and H). However the TT domain of *Lhx6* was gone (black arrowheads, G and H). E, F: The premamillary domain of *Lef1* was reduced (open arrowheads), as was the prethalamic domain (black arrowheads), and the anterior domain was absent in the mutants (yellow arrowhead). I, J: The thalamic eminence was unaffected (open arrowheads), as were the orexigenic neuron precursors labeled by *Lhx9*, although this domain was ventralized (black arrowheads). K-N: The PVN was unaffected in the mutant (black arrowheads). Markers encompassing the supramamillary and mamillary were reduced in the mutants (open arrowheads; K-P). The *Otp* domain of the anterior hypothalamus was unaffected (red arrowheads, K and L). The ZLI was unaffected (red arrowheads, M, N; black arrowheads O-R), as was the basal plate domain of Shh (open arrowheads; Q, R). The pituitary was displaced into the hypothalamic neuroepithelium (red arrowheads, S and T). *Pomc* in the arcuate was expanded into the posterior domain (black arrowheads, S and T). U: Schematic showing wild-type anatomy of the hypothalamus and prethalamus.

We next examined expression of *Otp* and *Sim1*, which both have discrete expression domains in posterior and anterior hypothalamus (Shimogori et al., 2010). In the case of *Otp* -- which is expressed in PvN, ArcN and PMN – we observed a substantial reduction in the size of the PMN expression domain (open arrowhead), a modest reduction in the size of the PvN domain (black arrowhead), and no change in the size of the ArcN domain (red arrowhead) (Fig. 3K-L, U). Likewise, a dramatic reduction in the size of the MM domain of *Sim1* was observed (open arrowhead), the PvN domain of Sim1 was moderately reduced (black arrowhead), and the ZLI domain was unaffected (red arrowhead) (Fig. 3M-N, U). A similar reduction in the size of the MM and SMN domain of *Pitx2* expression was observed (open arrowhead), while the ZLI domain was also unaffected (black arrowhead) (Fig. 3O-P, U). The posterior basal plate domain of Shh showed reduced expression (open arrowhead), while Shh expression in the ZLI (black arrowhead) and in the pituitary (red arrowhead) was not affected (Fig. 3Q-R, U). Finally, like in *Nkx2.1-Cre;Ctnnb1*^*lox/lox*^ mice, we observed a posterior expansion and dispersion of *Pomc*-expressing ArcN cells (Fig. 3S-U, Fig S3V-X).

### 3.3 Expression of constitutively active*Ctnnb1* in*Foxd1*-positive prethalamic neuroepithelium blocks expression of*Arx* and *Isl1*, but does not affect markers of prethalamic progenitors

We next investigated whether gain of function of *Ctnnb1* could produce phenotypes opposite to these seen following *Cttnb*1 loss of function. To achieve this, we generated *Foxd1-Cre;Ctnnb1*^*ex3*/+^ mice, in which Cre-dependent removal of an autoinhibitory domain in exon 3 of the *Ctnnb1* gene generates a constitutively active mutant form of the protein (Harada et al., 1999). By E12.5, *Foxd1-Cre;Ctnnb1*^*ex3*/+^ mice showed strong induction of *Lef1* expression throughout the prethalamus and hypothalamus, but not the EmThal, indicating robust and widespread induction of canonical Wnt signaling compared to littermate controls (Fig. 4A-B). Examination of other markers in constitutively active *Ctnnb1* mutants revealed a complex phenotype that does not directly mirror that observed in the loss of function mutants. *Foxd1-Cre;Ctnnb1*^*ex3*/+^ mice showed reduced expression of a number of prethalamic markers including *Arx* (black arrowhead) (Fig. 4C-D, U; Fig. S4A-F), *Isl1* (black arrowhead) (Fig. 4E-F, U; Fig. S4G-L), and *Lhx1* (red arrowhead) (Fig. 4G-H, U; Fig. S4M-R). Other markers of prethalamic progenitors, however, displayed only moderately reduced expression or no obvious change, including *Gsh2* (black arrowhead) (Fig. 4I-j, U) and *Fgf15* (black arrowhead) (Fig. 4K-L, U), indicating that general prethalamic identity was maintained in these mutants. Consistent with the fact that no induction of *Lef*1 expression was observed in EmThal, *Lhx1* expression in this structure was maintained (yellow and red arrowheads) (Fig. 4G-H, U; Fig. S4M-R).

**Figure 4:**
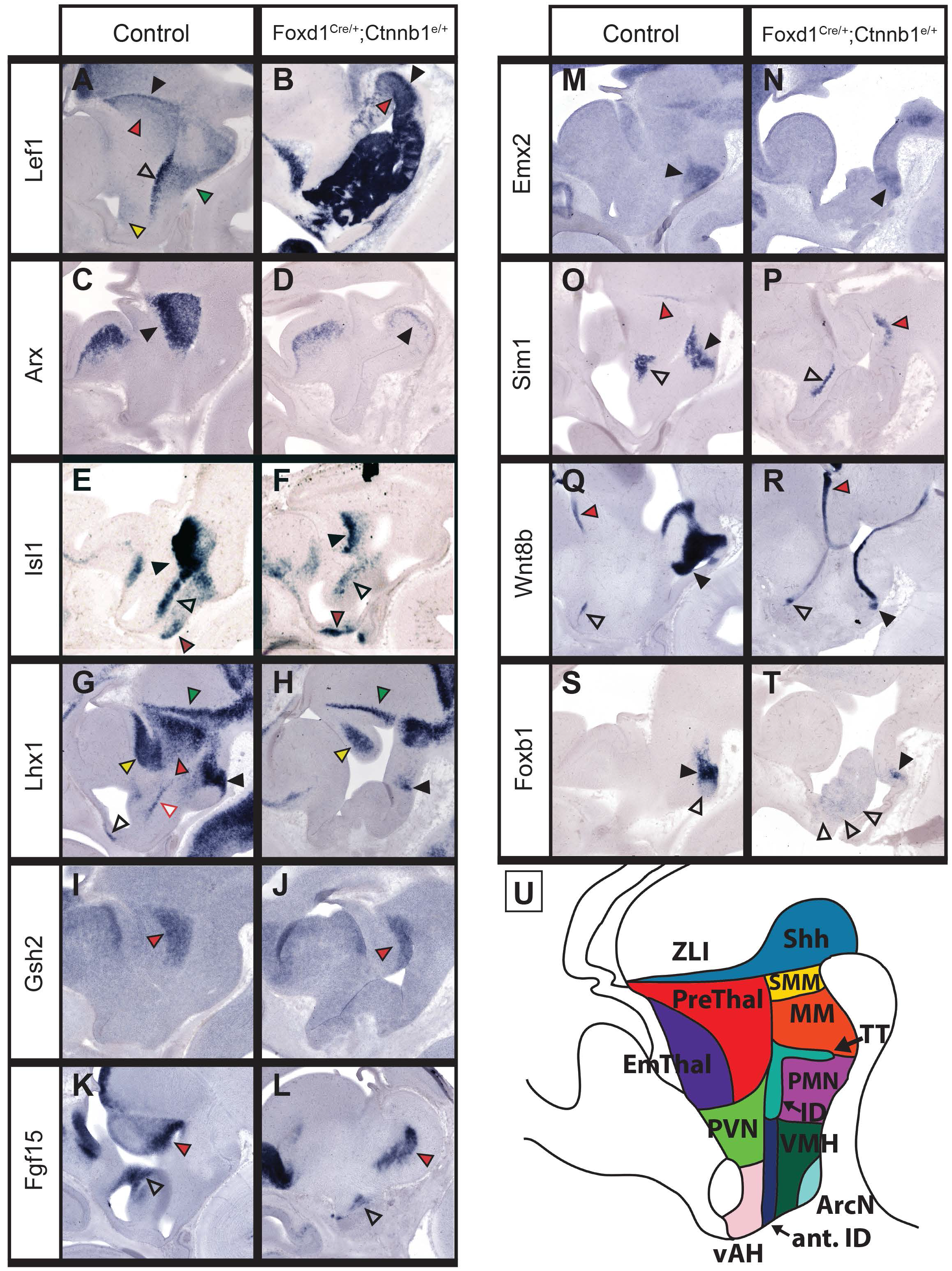
Constitutively active β-catenin results in disrupted patterning and development of the hypothalamus and prethalamus. A-T: *In situ* hybridization on E12.5 control (*Foxd1-Cre-*negative*;Ctnnb1*^*ex3*/+^) and *Foxd1-Cre;Ctnnb1*^*ex3*/+^ sagittal sections. A,B: *Lef1* was expanded throughout the hypothalamus, indicating a successful upregulation in canonical Wnt signaling. (Red arrowheads=prethalamus, black arrowheads=ZLI adjacent, green arrowhead=premamillary, open arrowhead=ID, yellow arrowhead=ventral AH). C-F: Black arrowheads indicate the prethalamus, which was absent except for a tiny sliver in D, and reduced but still present in F. Expression of the stripe marker *Isl1* was also reduced in the hypothalamus (open arrowheads), but not the ventral AH (red arrowheads) (E, F). G-L: *Lhx1* expression was completely lost in the prethalamus, but *Gsh2* and *Fgf15* expression were preserved (red arrowheads). *Lhx1* expression was also lost in the ID (red outlined arrowhead), anterior ID (open arrowhead) and heavily reduced in the mamillary (black arrowhead) but preserved in the thalamic eminence (yellow arrowhead) and ZLI (green arrowhead). *Fgf15* was also preserved in the dorsal anterior hypothalamus (open arrowheads; K, L). M-T: There were differential effects on mamillary markers in the mutant. *Emx*2 was largely preserved, while *Sim1* was completely absent (black arrowheads; M-P). The nuclear domains of *Wnt8b* and *Foxb1* were significantly reduced (black arrowheads; Q-T), although there was a migratory population of *Foxb1*-expressing cells that were expanded into the anterior domain in the GOF mutants (open arrowheads; S, T). *Sim1* was preserved in the PVN (open arrowheads) and ZLI (red arrowheads) (O, P), while the anterior domain (open arrowhead) and thalamic eminence domain (red arrowhead) of *Wnt8b* expression was similarly unaffected (Q, R). U: Schematic showing wild-type anatomy of the hypothalamus and prethalamus.

### 3.4 Expression of constitutively active*Ctnnb1* in*Foxd1*-positive hypothalamic neuroepithelium leads to expansion of a subset of posterior and premamillary markers, and a loss of expression of GABAergic neural precursor markers

Hypothalamic expression of *Arx*, *Isl1* and *Lhx1* was also severely disrupted. *Arx* expression in ID and TT was lost (open arrowhead) (Fig. 4C-D, U; Fig. S4A-F), while *Isl1* expression was greatly reduced in all hypothalamic regions (open arrowhead) except for ArcN (red arrowhead) (Fig. 4E-F, U; Fig. S4G-L). *Lhx1* expression was lost in the anterior ID, and substantially reduced in the posteriorly-located MM (black arrowhead) (Fig. 4E-F, U; Fig. S4G-L). Other markers of MM were also reduced, including *Emx2* (black arrowhead) (Fig. 4M-N, U), *Sim1* (Fig. 4O-P, U), *Wnt8b* (Fig. 4Q-R, U) and *Foxb1* (Fig. 4S-T, U). Although the nuclear MM expression domain of *Foxb1* was reduced, we observed widespread ectopic *Foxb1* expression in more anterior hypothalamic regions (open arrowhead), suggesting that these cells may have dispersed from the MM (Fig. 4S-T, U). Interestingly, we also observed either expanded or unchanged expression of other MM and SMN markers, such as *Lhx5* (open arrowhead) (Fig. 5A-B, U) and *Pitx2* (Fig. 5C-D, U; Fig. S4S-X).

**Figure 5:**
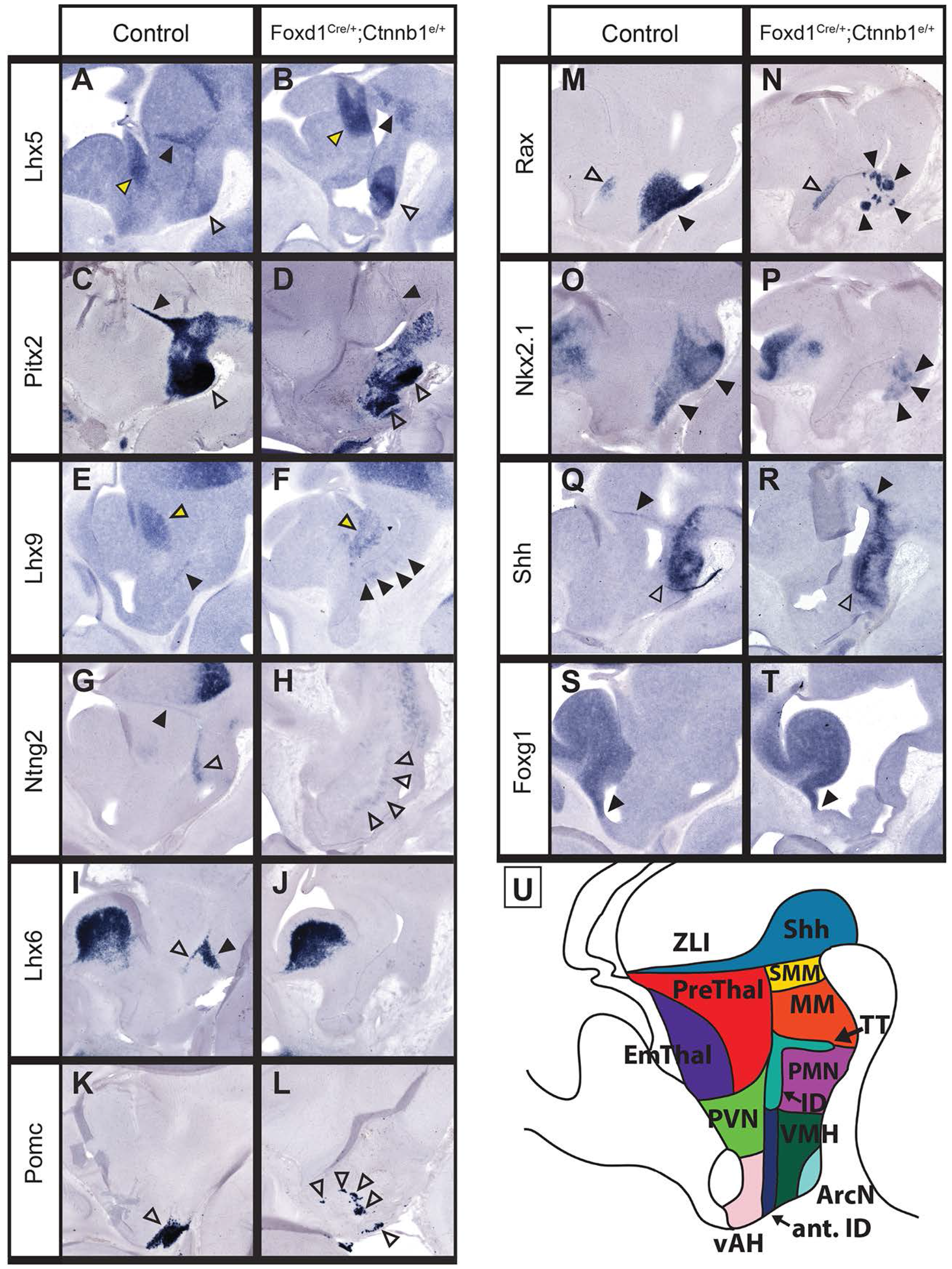
Constitutively active beta-catenin results in disrupted patterning and development of the hypothalamus and prethalamus. A-T: *In situ* hybridization on E12.5 control (*Foxd1-Cre-negative*;*Ctnnb1*^*ex3*/+^) and *Foxd1-Cre;Ctnnb1*^*ex3*/+^sagittal sections. A-D: Two markers of the posterior hypothalamus were expanded in the mutants: *Lhx5* in the mamillary (open arrowheads; A, B) and *Pitx2* in the SMN and MM (open arrowheads; C, D). ZLI expression was maintained for both of these markers (black arrowheads; A-D). Markers of the thalamic eminence were also preserved (yellow arrowheads; A, B, E, F). *Lhx9*-expressing orexigenic neurons were expanded in the GOF mutant (black arrowheads, E, F). G-H: The dorsal PMN expression of *Ntng2* was also expanded (open arrowheads). I-J: *Lhx6* was completely absent from the ID (open arrowhead) and TT (black arrowhead). K-L: The arcuate (open arrowheads) was severely disrupted in the GOF mutant. M-P: In the mutant, ventral stripe markers were reduced to their posterior domain and were extremely mosaic (black arrowheads) and expression of *Rax* within the PVN was unaffected (open arrowheads; M, N). Q-R: Expression of *Shh* was unaffected in the supramamillary/mamillary region (open arrowheads) and ZLI (black arrowheads). S-T: There was no shift in the location of the DTJ (black arrowheads). U: Schematic showing wild-type anatomy of the hypothalamus and prethalamus.

We also observed a complex phenotype in the anterodorsal hypothalamus. *Sim1* expression in the anterior PvN was substantially reduced (open arrow), although expression in the ZLI was maintained (Fig. 4O-P, U). Anterior hypothalamic domains of *Fgf15* (open arrowhead) (black arrowhead) (Fig. 4K-L) and *Wnt8b* (open arrowhead) (Fig. 4Q-R) were reduced and maintained, respectively. *Lhx9*, which marks precursors of orexinergic neurons (black arrowhead) (Fig. 5E-F, U; Fig. S5A-F) showed dispersed expression in more anterior hypothalamic structures similar to the phenotype seen with *Foxb1*. However, the EmThal expression domains of both *Lhx5* and *Lhx9* (Fig. 5A-B, 5E-F, U; Fig. S4S-X) were unaffected, like that of *Lhx1*.

*Ntng2*, which is expressed in PMH where *Lef1* levels are normally high (Shimogori et al., 2010), showed an expanded and anteriorly extended domain of hypothalamic expression (open arrowhead) (Fig. 5G-H, U; Fig. S5G-L). Expression of *Bsx*, which is detected in both PMN (black arrowhead) and ArcN (open arrowhead), was maintained in PMN but lost in ArcN (open arrowhead) (Fig. S5M-R). In addition to the loss of *Arx* and *Lhx1* in the ID (open arrowhead) and TT (black arrowhead), *Lhx6* was also lost in these structures in mutant animals (Fig. 5I-j, U; Fig. S5S-X). Expression of the ArcN marker *Pomc* was disrupted, including a reduced nuclear domain and ectopic domains of expression detected throughout the tuberal hypothalamus (Fig. 5I-j, U; Fig. S6A-F).

### 3.4 Expression of constitutively active*Ctnnb1* in*Foxd1*-positive hypothalamic neuroepithelium leads to expansion of a subset of posterior and premamillary markers, and blocks expression of markers of GABAergic precursors

These widespread changes in hypothalamic gene expression raised the question of whether expression of constitutively active *Ctnnb1* disrupted expression of genes required to maintain hypothalamic regional identity. We observed a substantial reduction and dispersal of *Rax*-positive cells in tuberal hypothalamus (black arrowhead), although expression of *Rax* in PvN progenitors was maintained (open arrowhead) (Fig. 5M-N, U; Fig. S6G-L). Expression of *Nkx2.1* was also reduced in tuberal hypothalamus, and likewise showed disrupted and dispersed expression in posterior hypothalamus (Fig. 5O-P). However, expression of Shh was maintained in the basal plate domain of hypothalamus (Fig. 5Q-R), and no change in the telencephalic expression domain of *Foxg1* was observed (Fig. 5S-T). These findings suggest that while overall hypothalamic identity is maintained in *Foxd1-Cre;Ctnnb1*^*ex3*/+^ mice, substantial changes in a subset of broadly expressed hypothalamic markers are observed.

### 3.5 Constitutively active *Ctnnb1* induces hyperproliferation of hypothalamic progenitors

To investigate the cellular mechanisms mediating the changes in gene expression detected in both *Foxd1-Cre;Ctnnb1*^*lox/lox*^ and *Foxd1-Cre;Ctnnb1*^*ex3*/+^ mice, we investigated changes in cell proliferation and death at E12.5. We observed a substantial increase in the thickness of both prethalamic and hypothalamic DAPI-positive neuroepithelium in *Foxd1-Cre;Ctnnb1*^*ex3*/+^ mice (Fig. S7A-D). EdU labeling at E12.5 revealed no significant change in proliferation in *Foxd1-Cre;Ctnnb1*^*lox/lox*^ (Fig. S7D-E), but a substantial increase in *Foxd1-Cre;Ctnnb1*^*ex3*/+^ mice (Fig. S7C). The increase in thickness and overall disorganization of the prethalamic and hypothalamic neuroepithelium is particularly clear in DTI imaging analysis of E12.5 *Foxd1-Cre;Ctnnb1*^*ex3*/+^ embryos (Fig. 6). In contrast to control embryos, where the ventricular margin of the neuroepithelium was smooth and radial glial fibers are aligned perpendicular to the ventricles (Fig. 6c), *Foxd1-Cre;Ctnnb1*^*ex3*/+^ embryos showed incipient foliation of hypothalamic and prethalamic neuroepithelium, along with corresponding disorganization of radial glial fibers (Fig. 6c). This massive disruption in hypothalamic development may contribute to the early death of *Foxd1-Cre;Ctnnb1*^*ex3*/+^ embryos, which do not survive past E15.

**Figure 6:**
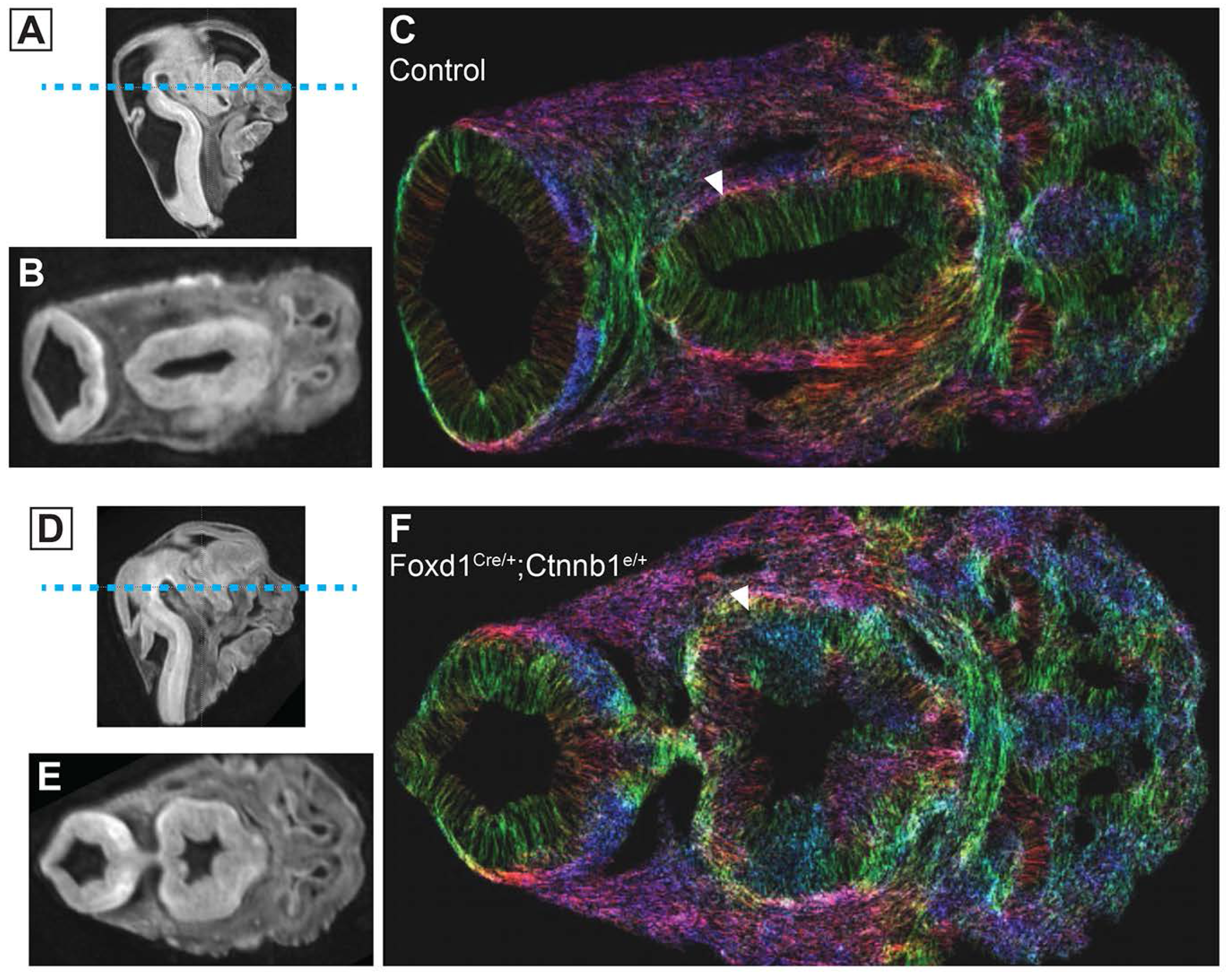
Hyperplastic hypothalamic neuroepithelium is observed in constitutively active beta-catenin mutants. A-F: MRI on *Ctnnb1*^*ex3*/+^ (A-C) and *Foxd1-Cre;Ctnnb1*^*ex3*/+^ (D-F) E12.5 embryos. A, D: diffusion-weighted MR images showing the gross neuroanatomy of the E12.5 embryos. Dashed line indicates the level at which the horizontal sections in B, C, E and F were taken. B, E: horizontal diffusion-weighted MR images at the level indicated in A and D. C and F: In the tract-density images, Colored streamlines indicate the orientation of the fiber tracts, which can either represent axons or radial glia, but in this case most likely represents radial glia. The hypothalamic neuroepithelium was significantly disrupted in the GOF mutant.

### 3.6 Gain and loss of function mutations in hypothalamic*Ctnnb1* both disrupt pituitary development

Finally, although neither the *Nkx2.1-Cre* nor the *Foxd1-Cre* lines are active in pituitary, we observed severe disruption of pituitary development following both loss and gain of function of *Ctnnb1* in hypothalamic neuroepithelium. At E12.5 in both *Foxd1-Cre;Ctnnb1*^*lox/lox*^ and *Nkx2.1-Cre;Ctnnb1*^*lox/lox*^ embryos, we observed that the anterior pituitary intruded into the tuberal hypothalamus, and that no pituitary stalk was observed (Fig. 7A-C). In contrast, in *Foxd1-Cre;Ctnnb1*^*ex3*/+^ embryos, the pituitary anlagen remained attached to the oral ectoderm (Fig. 7D-F). Furthermore, pituitary expression of both *Pitx2* and *Pomc* was substantially increased in *Foxd1-Cre;Ctnnb1*^*ex3*/+^ embryos.

**Figure 7:**
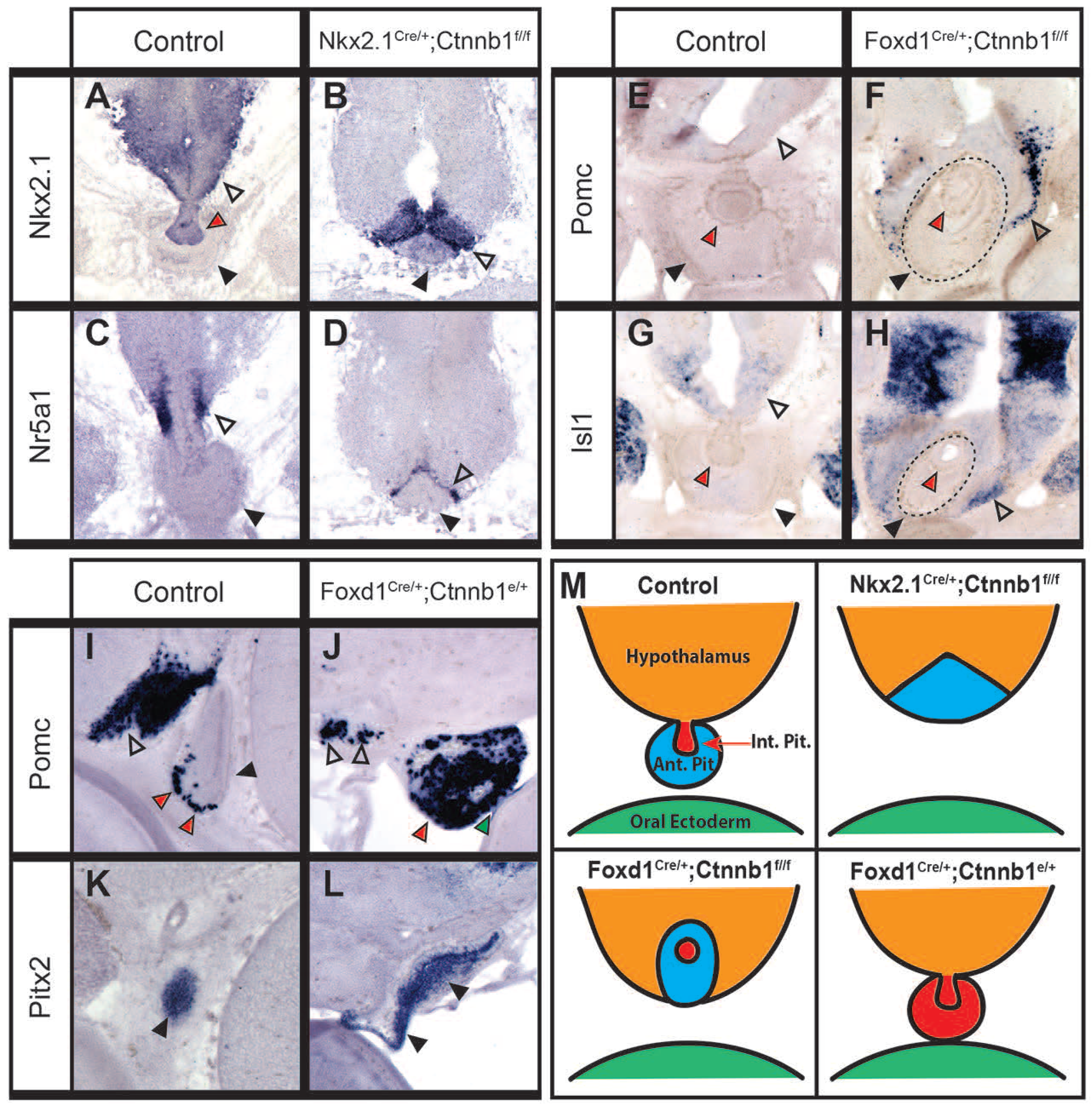
The specification and migration of the pituitary is disrupted by gain and loss of function of beta-catenin. A-D: *In situ* hybridization on E12.5 control (*Cre*-negative, either *Ctnnb1*^*lox/lox*^ or *Ctnnb1^lox/+^*) and *Nkx2.1-Cre;Ctnnb1*^*lox/lox*^ sagittal sections. A-B: The hypothalamic expression of *Nkx2.1* and *Nr5a1* (open arrowheads) was displaced into a wedge shape by the invading pituitary (black arrowheads) in the mutant. *Nkx2.1* is also expressed in the anterior pituitary (red arrowhead; A): but this domain could not be seen in the mutant. E-H: *In situ* hybridization on E12.5 control (*Cre*-negative, either *Ctnnb1*^*lox/lox*^ or *Ctnnb1^lox/+^*) and *Foxd1-Cre;Ctnnb1*^*lox/lox*^ coronal sections. The invading pituitary is demarcated with a dashed circle -- in this case, the pituitary maintained its shape (black arrowhead: pituitary; red arrowhead: anterior pituitary). *Pomc* and *Isl1* expression in the hypothalamus (open arrowheads) was displaced by the invading pituitary. I-L: *In situ* hybridization on E12.5 control (*Foxd1-Cre-*negative;*Ctnnb1*^*ex3*/+^) and *Foxd1-Cre;Ctnnb1*^*ex3*/+^sagittal sections. In constitutively active β-catenin mutant embryos, the pituitary was enlarged and had expanded expression of the anterior pituitary markers *Pomc* (red arrowheads, I, J) and *Pitx2* (black arrowheads, K, L). In at least one mutant, the pituitary remained stuck to the oral ectoderm (junction is indicated by green arrowhead in J). Open arrowheads in I, J indicate arcuate expression of *Pomc*. M: Schematic showing wild-type pituitary anatomy and the pituitary phenotypes observed in each mutant line examined.

**Figure 8:**
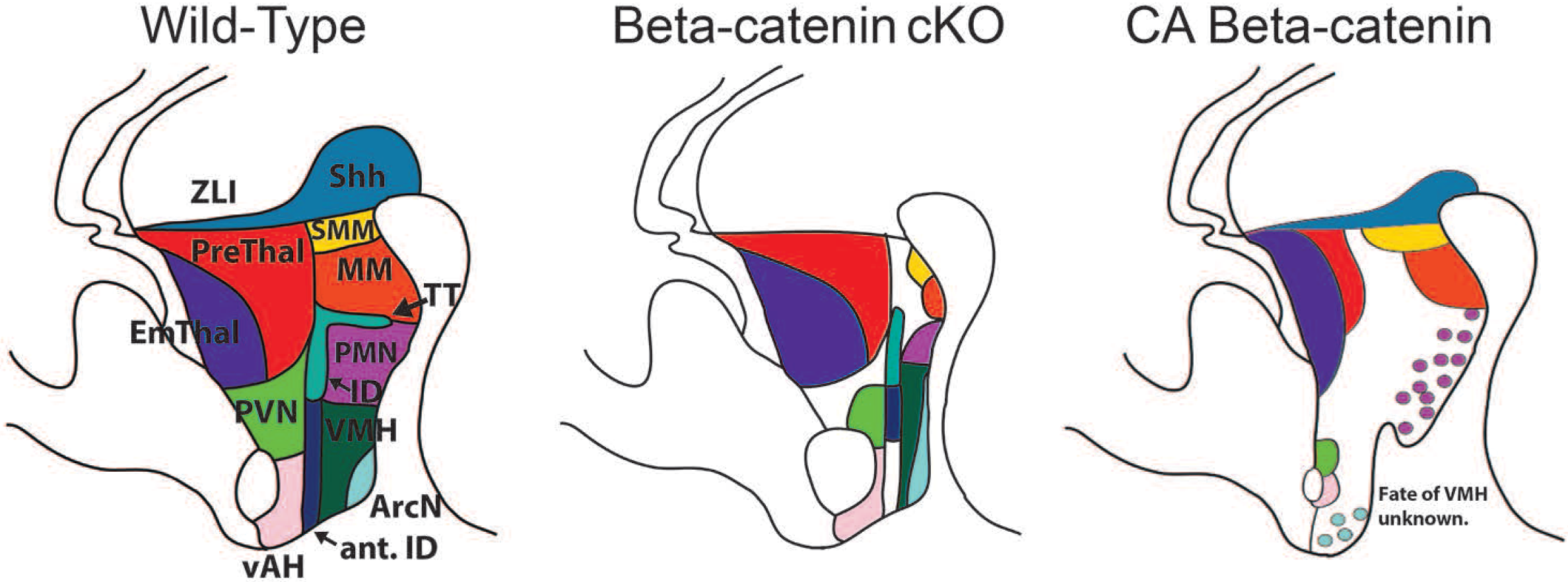
Schematic demonstrating the developmental changes observed in following gain and loss of function of beta-catenin. In LOF mutants, the hypothalamus was anteriorized, and thinned resulting in more dorsal structures shifting ventrally. The prethalamus was unaffected. In GOF mutants, the phenotype was much more complex, with some posterior markers expanded but others lost. For the most part, anterior markers were preserved but reduced and often mosaic. The prethalamus was smaller in size and missing expression of some, but not all markers. The fate of the VMH is unknown in these mutants

## 4 Discussion

This study demonstrates a central and complex role for canonical Wnt signaling in regulating the patterning and differentiation of mouse prethalamus and hypothalamus. Previous studies have demonstrated that canonical Wnt signaling confers posterior identity during neural development. During early stages of neural development, inhibition of Wnt signaling by Wnt antagonists secreted from the prechordal plate is necessary for forebrain specification (Shinya et al., 2000; Dorsky et al., 2003). Likewise, during early stages of thalamic development, Wnt signaling drives differentiation of the caudally located sensory thalamus, while repressing development of more rostral prethalamus (Bluske et al., 2012; Peng and Westerfield, 2006; Hagemann and Scholpp, 2012).

In this study, which examines the role of canonical Wnt signaling at later stages of prethalamic and hypothalamic development, we also observe phenotypes which are consistent with this pathway playing an important role in regulating anteroposterior patterning in hypothalamus. In both *Foxd1-Cre;Ctnnb1*^*lox/lox*^ and *Nkx2.1-Cre;Ctnnb1*^*lox/lox*^ loss of function mutants, we observe a decrease in expression of posterior hypothalamic markers, along with a corresponding expansion of anterior markers into posterior hypothalamus. In the hypothalamus of *Foxd1-Cre;Ctnnb1*^*ex3*/+^ gain of function mutations, expansion of a subset of posterior and premamillary markers is observed, along with a reduction or disruption in expression of a subset of anterior hypothalamic markers. In prethalamus, a gain of function mutation in *Ctnnb1* suppresses expression of GABAergic neuronal markers, while loss of function mutations lead to a much more modest reduction in expression of these markers. These data are consistent with a model in which canonical Wnt signaling acts as a posteriorizing signal in hypothalamic neuroepithelium, while acting to suppress expression of prethalamic markers.

A closer analysis of these results, however, reveals the role of canonical Wnt signaling to be substantially more complex, particularly upon closer examination of the effects of the gain of function mutation. While some posterior hypothalamic markers such as *Pitx2* and *Lhx5* were expanded, other posterior markers such as *Emx2*, and *Lhx1* were reduced. Genes selectively expressed in the *Lef1*-positive premamillary hypothalamus, such as *Lhx9* and *Ntng2*, showed expanded expression, as expected, and consistent with studies in zebrafish (Lee et al., 2006; Xie et al., 2017). However expression of the TT marker *Lhx6* – which is expressed in GABAergic neurons in the premamillary region -- which shows high expression of *Lef1* – was completely lost in these mutants. GABAergic markers expressed in the hypothalamic ID – such as *Lhx1, Lhx6* and *Arx* – were also lost. In contrast, other anterior hypothalamic markers expressed in mitotic progenitors or glutamatergic cells -- such as *Pomc*, *Fgf15*, *Otp*, *Sim1*, and the anterior hypothalamic domains of *Rax* and *Wnt8b* – were preserved, although often showed reduced or disrupted expression. Moreover, markers of prethalamic progenitors such as *Gsh2* and *Fgf15* were essentially unaltered in *Ctnnb1* gain of function mutations, implying that prethalamic regional identity was preserved.

These findings show some major differences, but also important parallels, with previous studies on the role of canonical Wnt signaling in diencephalic development. Previous studies on the canonical Wnt effector *Lef1* in developing zebrafish hypothalamus, demonstrated that *Lef1* loss of function did not affect overall hypothalamic patterning, but blocked expression of *Isl1* and *Dlx2a* in premamillary hypothalamus (Lee et al., 2006). While we do observe a reduction in hypothalamic *Arx* and *Isl1* expression in *Ctnnb1* loss of function mutations, we also observe a loss of other posterior hypothalamic markers. Moreover, while zebrafish *Lef1* mutants do exhibit a smaller posterior hypothalamus, this results from defects in postembryonic neurogenesis (Wang et al., 2012), in contrast to the defects seen in mouse *Ctnnb1* mutants.

Studies of early thalamic development have demonstrated that canonical Wnt signaling promotes expression of sensory thalamic markers while suppressing expression of prethalamic markers (Bluske et al., 2012; Braun et al., 2003; Martinez-Ferre et al., 2013; Mattes et al., 2012). We observed that *Lhx9* and *Ntng2*, along with *Lef1* itself, were prominently expressed in both premamillary hypothalamus and sensory thalamus, and that these genes showed similar changes in response to gain and loss of function. This implies that at least a subset of the core beta-catenin-regulated transcription network that is involved in Wnt-mediated patterning of the sensory thalamus may also be used in patterning of premamillary hypothalamus.

Furthermore, we observed that genes that selectively label GABAergic neural precursors in prethalamus -- such as *Isl1*, *Arx* and *Lhx1* – also showed drastically reduced or absent expression in these animals. Strikingly, hypothalamic *Arx* expression was also lost, as was *Lhx1* expression in GABAergic neuronal precursors of the ID, while both *Lhx1* expression in glutamatergic mamillary precursors and hypothalamic *Isl1* expression, which is essential for specification of multiple neural subtypes in ArcN (Lee et al., 2016; Nasif et al., 2015), were both dramatically reduced (Shimogori et al., 2010). In sharp contrast, however, expression of the prethalamic progenitor markers *Fgf15* and *Gsh2* was unaffected (Shimogori et al., 2010). Moreover, sensory thalamic markers such as *Lhx9* and *Ntng2* were not induced in prethalamic neuroepithelium. This demonstrates that constitutively high beta-catenin activity does not in fact affect initial specification of prethalamic neuroepithelium, at least when induced after the onset of *Foxd1* expression, but instead potently inhibits generation and/or differentiation of GABAergic neurons of both the prethalamus and hypothalamus.

In addition to these effects on patterning and neuronal differentiation, we observed a massive increase in cell proliferation in *Ctnnb1* gain of function mutants that was not observed in previous studies, which used *Olig3-Cre* to induce expression of constitutively active beta-catenin (Bluske et al., 2012). Although many markers of prethalamic and hypothalamic neurons are lost in the *Foxd1-Cre;Ctnnb1*^*ex3*/+^ mutant, these animals showed a dramatic increase in proliferation and a large number of ectopic *Lef1*-positive cells. The precise identity of these cells is unclear. *Lef1*-expressing cells were increased to a much greater extent than were markers such as *Lhx9* and *Ntgn2*, implying that it is unlikely that these cells have been respecified to a premamillary identity. Most likely, these cells are simply arrested in an immature state, as suggested by studies in other CNS cell types (Wrobel et al., 2007;Joksimovic et al., 2009). Future studies, particularly those using single-cell RNA-Seq analysis (Picelli, 2017), can resolve this question in an unbiased and high-throughput manner. In contrast, *Ctnnb1* loss of function mutants did not show any obvious changes in proliferation or cell death at E12.5.

Finally, genetic manipulation of *Ctnnb1* unexpectedly led to region-specific disruption of cell organization in hypothalamus. Both gain and loss of function mutants of *Ctnnb1* led to *Pomc*-positive cells being dispersed throughout the anterior and tuberal hypothalamus, and lost the compact nuclear morphology seen in wildtype animals. Moreover, in *Ctnnb1* gain of function mutants, *Foxb1*-expressing cells of the MM, as well as *Lhx9* and *Ntng2*-expressing cells of the dorsal PMN, all showed broad dispersion throughout the posterior and tuberal hypothalamus. These phenotypes may be due to disrupted expression of genes controlling cell adhesion, resulting in disrupted nucleogenesis. A similar mechanism regulates cell distribution in the thalamus, where canonical Wnt signaling acts in developing thalamus to control *Pcdh10b* expression, and cells in the caudal and rostral thalamus intermingle when these domains express similar levels of *Pcdh10b* (Peukert et al., 2011).

Taken together, these findings show that canonical Wnt signaling regulates many different aspects of prethalamic and hypothalamic development, including regional patterning, neuronal differentiation, progenitor proliferation, and nucleogenesis. There are some clear similarities in the mechanism of action of this signaling pathway in both prethalamus and hypothalamus. Most notably, *Ctnnb1* gain of function mutants show a loss of expression of common GABAergic neuronal precursor markers such as *Arx* and *Lhx1*, while a much more modest reduction in expression of these markers is seen in loss of function mutations. Similar phenotypes are also seen for *Isl1*. The similarity in the prethalamic and hypothalamic phenotypes of these mutants is less surprising when one considers that these genes are expressed in contiguous and partially overlapping domains in both hypothalamus and prethalamus. *Arx*-positive GABAergic neurons of the prethalamus form a contiguous domain with hypothalamic ID and TT, and express a large number of common molecular markers (Shimogori et al., 2010). *Isl1*-expressing cells likewise comprise a contiguous, and partially overlapping, zone that includes both GABAergic cells in the prethalamus and a highly diverse assortment of neuronal subtypes in the hypothalamus (Lee et al., 2016; Nasif et al., 2015; Shimogori et al., 2010). These results are the latest in a series of genetic studies (MacDonald et al., 2013; Roy et al., 2013) that cannot readily be accounted for by the prosomere model of forebrain organization (Puelles and Rubenstein, 2015), and instead indicate that the hypothalamus and prethalamus form a common developmental unit, as predicted by traditional columnar models of brain organization (Swanson, 1992, 2012).

The source and identity of the Wnt ligand controlling beta-catenin activity in hypothalamus and prethalamus is still undetermined. It is likely that *Wnt8a/b* derived from mamillary hypothalamus is mediating the patterning of the hypothalamus, consistent with research in zebrafish (Kim et al., 2002; Lee et al., 2006). Although *Wnt8b*^−/−^ mice were reported to have no discernible defects in patterning in the diencephalon (Fotaki et al., 2010), this study only investigated expression of three diencephalic markers, one of which was *Isl1*, a gene whose expression domain was displaced ventrally in *Ctnnb*1 loss of function mutants, but did not otherwise display a shift in position or expression. It is likely that other Wnts are compensating for the loss of *Wnt8b* in regulating hypothalamic patterning, with the best candidate being *Wnt8a*, which is expressed in a similar pattern to *Wnt8b*. The selective defects in development of prethalamic and hypothalamic GABAergic neurons seen in our mutants may reflect a differential requirement for Wnt signaling in their development. Since this population selectively expresses Wnt7a and Wnt7b (Shimogori et al., 2010), it is possible that these factors may serve as a paracrine signal to control the differentiation of these cells.

Finally, one of the more surprising results of this study was the disruption of the pituitary in all mutant lines examined. Normally, the anterior pituitary forms from the oral ectoderm and migrates to meet the posterior pituitary ventral to the hypothalamus. In both loss of function *Ctnnb1* mutants, the migrating pituitary appeared to have failed to receive any ‘stop’ signal and was invading the hypothalamus itself. In the beta-catenin gain of function mutant, the opposite phenotype occurred. The pituitary failed to migrate at all, and remained stuck to the roof of the mouth. It also exhibited a fate change, as the *Pomc* and *Pitx2* expression domains were significantly expanded. The effects on pituitary development are non-cell autonomous, as the *Foxd1-Cre* is not expressed in the pituitary. Instead, this effect must have been mediated by changes in the hypothalamus, which secretes many important signaling molecules important for pituitary development, including FGFs and BMP7 (Potok et al., 2008). An apoptotic domain on the ventral side of Rathke’s pouch is necessary for separation from the oral ectoderm; this domain is most likely lacking in the *Ctnnb1* gain of function mutant, perhaps due to increased expression of prosurvival factors from the hypothalamus (Davis and Camper, 2007). Altered expression of BMPs and BMP antagonists such as noggin may be mediating the expanded marker domains, as this pathway is known to regulate *Pomc* expression in the pituitary (Davis and Camper, 2007). More research needs to be done to determine the cause of the migratory defect observed in hypothalamic-specific loss of function mutants of *Ctnnb1* and the precise cause of the fate shift seen in gain of function mutants.

## Acknowledgements

We thank Wendy Yap and Thomas Kim for comments on the manuscript. This work was supported by NIH R01DK108230, and a Hopkins Synergy Award (to S.B.)

## Author contributions

EAN conducted experiments and analyzed all data. DW and JZ performed DTI analysis. MMT provided *Ctnnb1*^*ex3*/+^ mice. EAN and SB wrote the manuscript, with input from all authors.

## Author information

The authors declare no competing financial interests.

## Abbreviations

ArcN: arcuate nucleus
EmThal: thalamic eminence
ID: intrahypothalamic diagonal
MM: mamillary hypothalamus
PMN: premamillary hypothalamus
PrThal: prethalamus
PvN: paraventricular nucleus
SMN: supramammilary hypothalamus
SCN: suprachiasmatic nucleus
VMH: ventromedial hypothalamus

